# Disentangling the role of poultry farms and wild birds in the spread of highly pathogenic avian influenza virus H5N8 in Europe

**DOI:** 10.1101/2021.10.22.465255

**Authors:** Claire Guinat, Cecilia Valenzuela Agui, Timothy G. Vaughan, Jérémie Scire, Anne Pohlmann, Christoph Staubach, Jacqueline King, Edyta Swieton, Adam Dan, Lenka Cernikova, Mariette F. Ducatez, Tanja Stadler

**Affiliations:** Department of Biosystems Science and Engineering, ETH Zurich, Switzerland; Swiss Institute of Bioinformatics (SIB), Switzerland; Friedrich-Loeffler-Institut, Germany; National Veterinary Research Institute, Poland; Danam Vet Molbiol, Koszeg, Hungary; State Veterinary Institute, Prague, Czech Republic; INRAE-ENVT, France

**Keywords:** phylodynamics, Europe, highly pathogenic avian influenza, H5N8, virus transmission and evolution, Bayesian inference

## Abstract

Recent outbreaks of highly pathogenic avian influenza H5N8 virus in Europe have caused severe damage to animal health, wildlife conservation and livestock economic sustainability. While epidemiological and phylogenetic studies have generated important clues about the virus spread in Europe, they remained opaque to the specific role of poultry farms and wild birds. Using a phylodynamic framework, we inferred the H5N8 virus transmission dynamics among poultry farms and wild birds in four severely affected countries and investigated drivers of spread between farms across borders during the 2016-17 epidemic. Based on existing genetic data, we showed that the virus was likely introduced into poultry farms during the autumn, in line with the timing of arrival of migratory wild birds. Then, transmission was mainly driven by farm-to-farm transmission in Germany, Hungary and Poland, suggesting that better understanding of how infected farms are connected in those countries would greatly help control efforts. In contrast, the epidemic was dominated by wild bird-to-farm transmission in Czech Republic, meaning that more sustainable prevention strategies should be developed to reduce virus exposure from wild birds. We inferred effective reproduction number *R*_e_ estimates among poultry farms and wild birds. We expect those estimates being useful to parameterize predictive models of virus spread aiming at optimising control strategies. None of the investigated predictors related to live poultry trade, poultry census and geographic proximity were identified as supportive predictors of the viral spread between farms across borders, suggesting that other drivers should be considered in future studies.

**Significance statement:** In winter 2016-17, Europe was severely hit by an unprecedented epidemic of highly pathogenic avian influenza (HPAI) H5N8 virus, causing significant impact on animal health, wildlife conservation and livestock economic sustainability. By applying phylodynamic tools to H5N8 sequence data collected from poultry farms and wild birds during the epidemic, we quantified how effectively the first infections were detected, how fast the virus spread, how many infections were missed and how many transmission events occurred at the wildlife-domestic interface. Also, we investigated predictors of the virus spread between farms across borders. These results are crucial to better understand the virus transmission dynamics, with the view to inform policy decision-making and reduce the impact of future epidemics of HPAI viruses.

## Introduction

Since the beginning of the 21^st^ century, the highly pathogenic avian influenza (HPAI) H5N8 virus (clade 2.3.4.4b) represents one of the most serious threats to animal health, wildlife conservation and livestock economic sustainability. In June 2016, the virus was detected in wild birds in regions of Central Asia (at the Ubsu-Nur and Qinghai lakes, known as migration stop-overs) and subsequently spread to other Asian countries and Europe (1). By the end of 2017, the virus had caused one of the most severe epidemic in Europe, in terms of number of poultry outbreaks, wild bird cases and affected countries (1). Most of poultry outbreaks occurred in France (37.8%), followed by Hungary (21.5%), Germany (8.5%), Poland (5.8%) and Czech Republic (3.9%) (1). While no human cases were observed, the control strategies that were implemented in the affected countries resulted in the culling of several million poultry, causing devastating socio-economic impacts for the poultry industry.

The emergence of H5N8 virus in Europe was likely attributable to infected migratory wild birds from Northern Eurasia, leading to occasional or multiple viral incursions into poultry farms (2–7). After emergence, farm-to-farm transmission was likely the main driver of the epidemic, with contact with infected poultry and contaminated fomites, such as vehicles or equipment, being a major risk factor for farm infection (1–3, 6, 8, 9). In a number of cases, high poultry density and substantial gaps in farm biosecurity were also identified as potential risk factors for farm infection (1, 3, 6, 10). The possibility of airborne transmission between poultry farms was also suggested, without being conclusively demonstrated (11, 12).

While such epidemiological and phylogenetic studies have generated important clues about the H5N8 virus transmission patterns in Europe, they remained opaque to the specific role of poultry farms and wild birds in the disease spread. In particular, understanding the viral transmission dynamics among these two subpopulations is crucial to determine which of these two has the greatest potential to drive the viral transmission during epidemics, which, in turn, represents critical information to better target control strategies. When appropriate pathogen genetic and epidemiological data is collected, phylodynamic methods can fill this critical gap (13–15). By fitting population dynamic models to genetic sequences collected during epidemics, these tools aim at quantifying disease transmission dynamics and have been particularly used to study the spread of infectious diseases in structured populations, be they stratified by time, species or geography (16–18). Importantly, birth-death model-based approaches (19) explicitly allow for the direct estimation of key epidemiological parameters, such as the effective reproduction number *R*_e_ (which captures the number of secondary infections generated at any time during an epidemic in a partially immune population) (20), while taking into account the sampling effort.

Using a phylodynamic framework, this study aimed at disentangling the role of poultry farms and wild birds in the spread of H5N8 in Europe during the 2016-2017 epidemic. We fitted a phylodynamic model with geographical and host structure to H5N8 genome sequences of the HA segment collected from both host types (190 from poultry farms and 130 from wild birds) in four severely affected European countries (Czech Republic, Germany, Hungary and Poland) to: (i) estimate the early patterns of virus spread, (ii) infer the number of unreported infections, (iii) provide *R*_e_ estimates, (iv) discriminate the number of new infections arising from local transmission versus importation events and (v) identify factors driving the virus spread between farms across borders.

## Results

We fitted a multi-type birth-death (MTBD) model to the aligned sequences (19, 21) to co-infer epidemiological parameters, along with the underlying structured phylogenetic trees (22) and epidemic trajectories (23) (see Materials and Methods). The model was structured according to the host type and geographical location, resulting into five demes: ‘poultry farms in Czech Republic’, ‘poultry farms in Germany’, ‘poultry farms in Hungary’, ‘poultry farms in Poland’ and ‘wild birds in the four countries’. This latter was assumed to represent the epidemic origin (24) and all transmission, become non-infectious and sampling processes were assumed to be deme-specific and constant through time (except for the within-deme *R*_e_ that was estimated across four-time intervals) (19, 21).

### Early patterns of H5N8 virus spread in Europe

The maximum clade credibility (MCC) tree reconstructed using the MTBD model is shown in Figure 1. Sequences from poultry farms of the same country for Germany, Hungary and Poland were generally clustered together in the tree, while sequences from poultry farms for Czech Republic were more scattered in the tree as were wild birds’ sequences. From the epidemic trajectories, we extracted the first transmission events and summarized them over time to inform the early patterns of virus spread. Figure 2 shows the temporal distribution of inferred dates of the first imported (i.e. from another deme), local (i.e. within-deme) and exported (i.e. to another deme) outbreak/case per deme together with the first officially reported outbreak/case for comparison (25). Overall, the inferred dates of the first imported poultry farm outbreak were before the date of the first officially reported outbreak in each deme, with a higher delay in Czech Republic deme (median: 115 days, 95% High Posterior Density (HPD): 70 – 158), i.e. approximately 16 weeks) compared to the others (from median: 21 days, 95% HPD: 5 – 54 to median: 46 days, 95% HPD: 5 – 106, i.e. approximately 3 to 7 weeks) (SI Appendix Table S1). The first events of local virus transmission and exportation also took place before the first poultry farm outbreak was officially reported in each deme and occurred very rapidly after the first virus introduction (SI Appendix Table S1).

**Figure 1.**
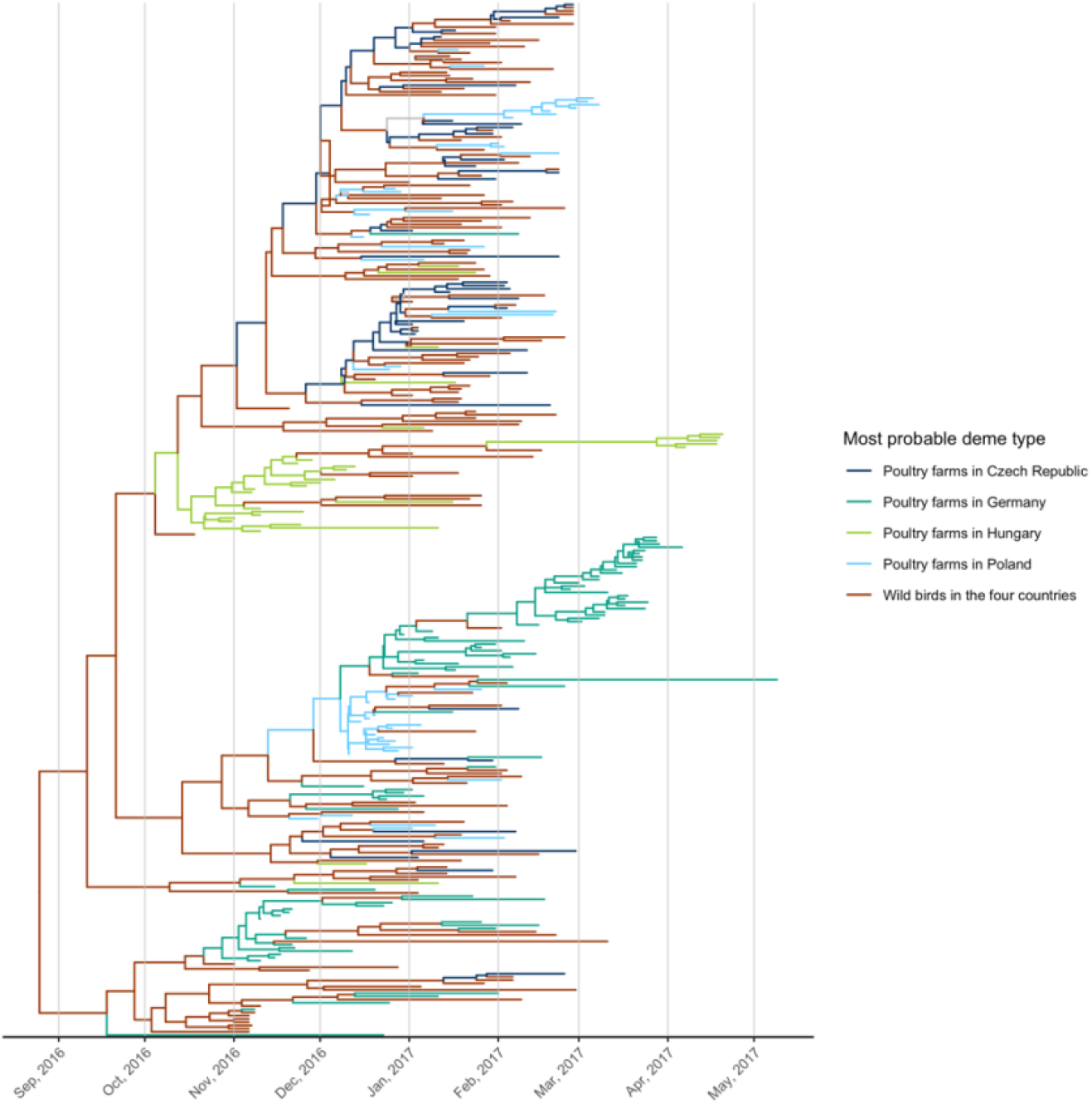
Time-scaled maximum clade credibility phylogenetic tree of the HA segment of HPAI H5N8 virus sequenced from poultry farms and wild birds during the 2016-2017 epidemic in Czech Republic, Germany, Hungary and Poland. The color of the tree branches shows the deme type with the highest probability (see legend) and uncertainty in deme type assignment is shown in grey. There is evidence for virus spread among neighbouring poultry farms illustrated by the presence of clusters of H5N8 sequences from poultry farms of the same country (mainly Germany, Hungary and Poland) in the tree, with the possibility of wild birds’ movements facilitating virus spread between poultry farms across countries, illustrated by the dispersal distribution of H5N8 sequences from wild birds.

**Figure 2.**
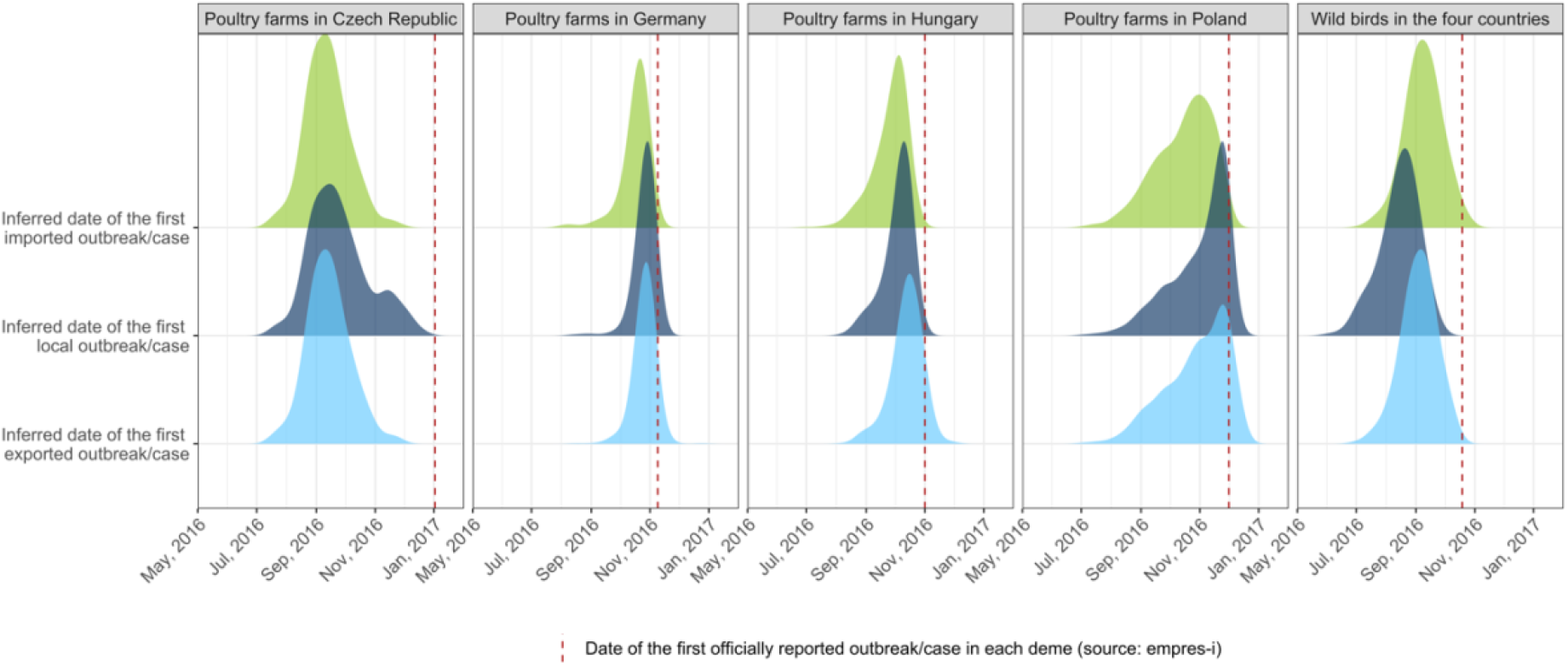
Temporal distribution of the inferred date of the first imported, local and exported outbreak/case per deme. The red dashed line represents the first officially reported outbreak/case per deme for comparison. In this graph, for each trajectory and each deme, we extracted the date of first imported, local and exported outbreak/case and summarized them over time in a probability density.

### Number of unreported H5N8 infections

From the epidemic trajectories, we extracted the number of poultry farms and wild birds that became non-infectious (following death, culling or recovery of the poultry flock/wild bird) and summarized it over time. This number was used to inform the number of outbreaks/cases that could have been missed during the epidemic. Figure 3 represents the temporal distribution of the inferred cumulative number of no-longer infectious outbreaks/cases per deme together with the cumulative number of officially reported outbreaks/cases for comparison (25). The cumulative number of officially reported poultry farm outbreaks (94 for Germany, 240 for Hungary and 65 for Poland) were within the inferred 95% HPD (median: 80, 95% HPD: 26-1196 for Germany, median: 401, 95% HPD: 142-1126 for Hungary and median: 96, 95% HPD: 34-398 for Poland). More discrepancies were observed for poultry farms in Czech Republic and wild birds in the countries, the cumulative number of officially reported outbreaks/cases (43 for Czech Republic and 372 for wild birds) being outside the inferred 95% HPD (median: 134, 95% HPD: 54 – 356 for Czech Republic and median: 4062, 95% HPD: 1138-14569 for wild birds).

**Figure 3.**
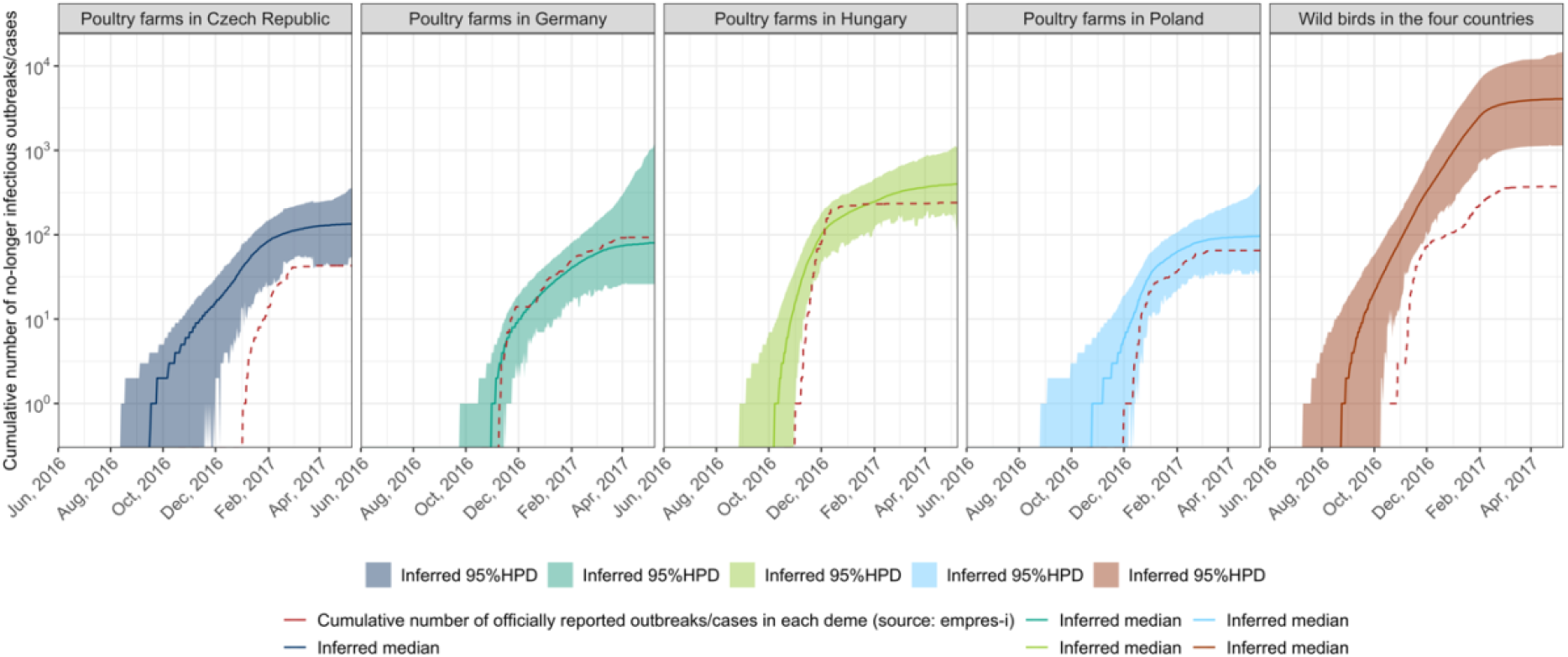
Temporal distribution of the inferred cumulative number of no longer infectious outbreaks/cases per deme. The solid line represents the median inferred, the colored areas represent the 95% HPD. The red dashed line represents the cumulative number of officially reported outbreaks/cases in log scale (25). In this graph, for each trajectory and each deme, we extracted the cumulative number of become non-infectious events and summarized them over time in log scale.

### Key epidemiological parameters of H5N8 virus spread

Figures 4A and 4B illustrate the posterior distributions for within-deme and between-deme *R*_e_ respectively, together with the prior (SI Appendix Table S2) for comparison. The within-deme *R*_e_ was estimated across four-time intervals, corresponding to the four phases of the epidemic (SI Appendix Figure S1). Note that only one sequence was available per poultry farm and per wild bird, meaning that the within-deme *R*_e_ represents the farm-to-farm/wild bird-to-wild bird virus transmission and the between-deme *R*_e_ represents the cross-species and cross-country virus transmission (i.e. farm-to-wild bird/wild bird-to-farm/farm-to-farm across countries). For most demes, the median within-deme *R*_e_ posteriors were above or close to 1 during the first time period, while a decrease was generally observed throughout the subsequent time periods (Figures 4A, SI Appendix Table S3). However, these *R*_e_ estimates slightly increased again during the fourth time period (Feb – May 2017) in Germany, Hungary and Poland. The highest median *R*_e_ estimates were observed between poultry farms in Hungary and between wild birds in the four countries during the first time period (Oct – Nov 2016). Overall, the between-deme *R*_e_ estimates were much lower than the within-deme *R*_e_ estimates, with extremely low values (median within the range of 10^−3^ – 10^−2^) for those representing farm-to-farm across countries and wild bird-to-farm transmission (Figure 4B, SI Appendix Table S3). Slightly higher values were found for those representing farm-to-wild bird transmission (median of 0.1 – 0.4). One exception was found for the between-deme *R*_e_ estimate representing farm-to-wild bird transmission in Czech Republic, with a median of 4.6 (95% HPD: 0.9-9.2). The infectious period was also inferred for each deme, with the highest median found for poultry farms in Czech Republic (median: 14 days, 95% HPD: 7 – 23) and wild birds in the four countries (median: 14 days, 95% HPD: 11 - 19) (SI Appendix Figure S2 and Table S3). The infectious period was slightly lower for poultry farms in Germany (median: 10, 95%HPD: 6 – 16), Hungary (median: 8 days, 95% HPD: 4 – 14) and Poland (median: 7 days, 95%HPD: 4 – 11).

**Figure 4.**
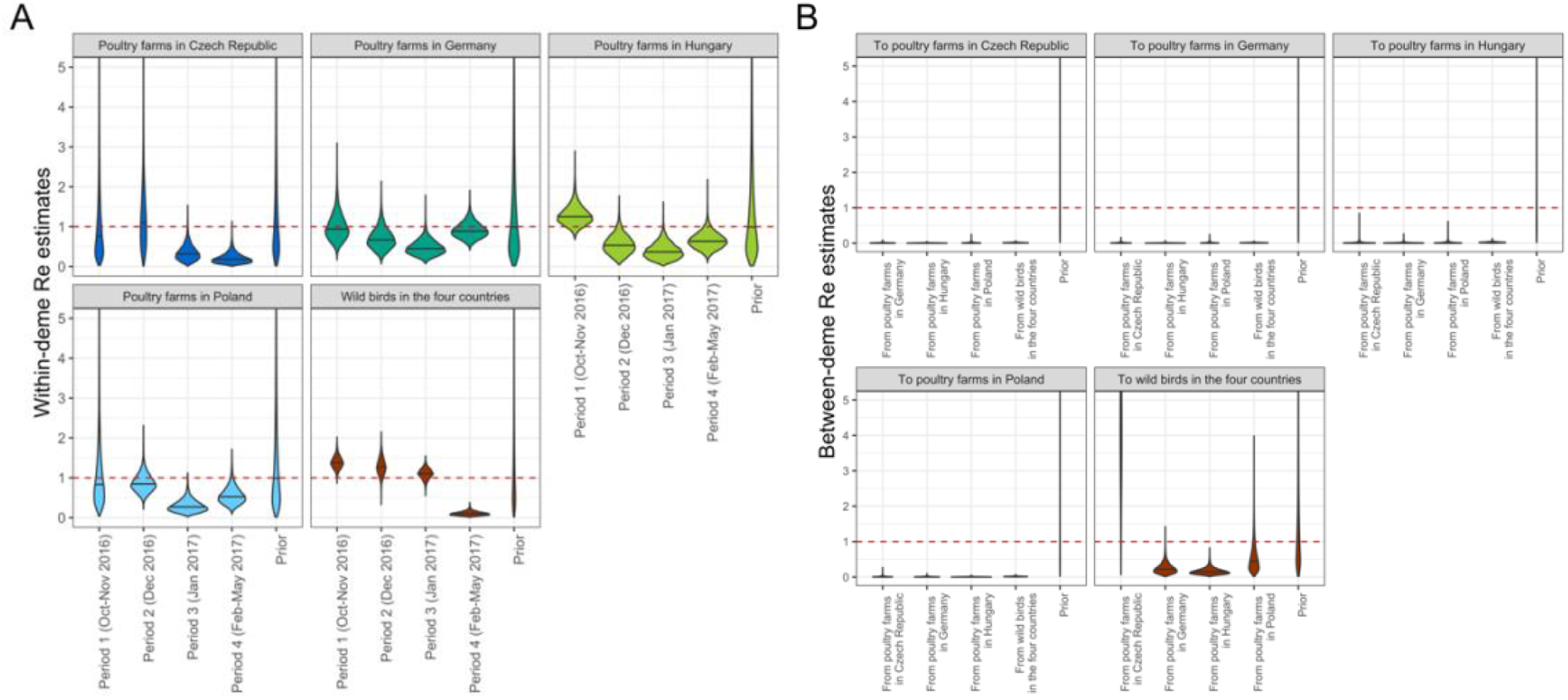
A) Posterior distributions for the within-deme *R*_e_ values across four-time intervals. Solid horizontal lines represent median values and the dashed red line represents the threshold between epidemic growth and fade out. B) Posterior distributions for the between-deme *R*_e_ values. Solid horizontal lines represent median values and the dashed red line represents the threshold between epidemic growth and fade out.

### Number of local H5N8 transmission versus importation events

From the epidemic trajectories, we can discriminate the number of poultry farm outbreaks and wild bird cases arising from local virus transmission (i.e. farm-to-farm/wild bird-to-wild bird) versus virus importation (i.e. farm-to-wild bird/wild bird-to-farm/farm-to-farm across countries) events. Figure 5 illustrates the temporal distribution of the inferred median number of local transmission and importation events per deme. In Germany, Hungary and Poland, the epidemic was dominated by local virus transmission events between poultry farms (median: 109, 95% HPD: 61 – 5750, median: 316, 95% HPD: 144 – 888 and median: 77, 95% HPD: 32 – 367, respectively) with an increase around March 2017, November 2016 and December 2016, respectively (SI Appendix Table S4). In Czech Republic, the epidemic was dominated by importations from wild birds (median: 115, 95% HPD: 57 – 230). For all countries, an increase in the number of importation events from wild birds were observed around January – February 2017. The epidemic in wild birds was also dominated by local transmission between wild birds (median: 3,075, 95% HPD: 807 – 8,575) and the highest number of importations were coming from poultry farms in Czech Republic (median: 972, 95% HPD: 77 – 5,569).

**Figure 5.**
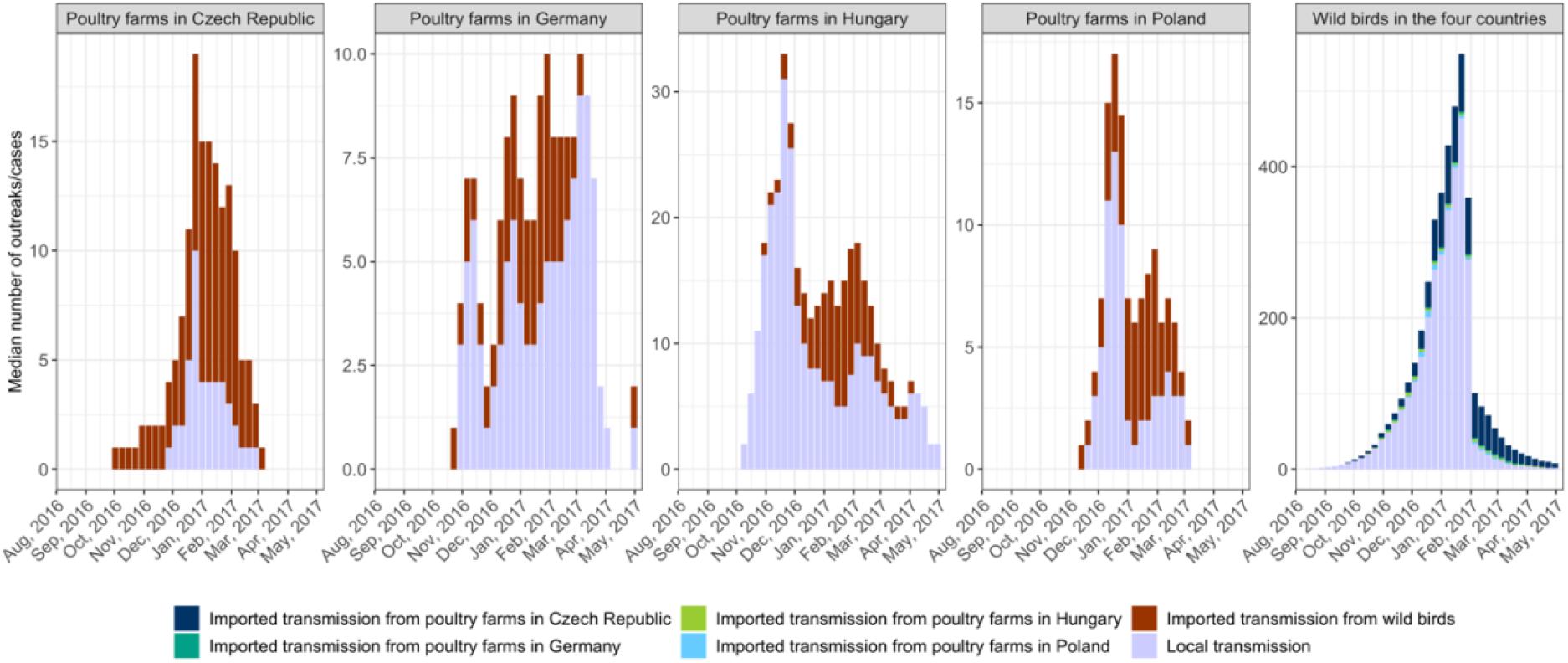
Temporal distribution of the inferred median number of poultry farm outbreaks and wild bird cases arising from local transmission and importation events per deme. In this graph, for each trajectory and each deme, we computed the median number of within-deme and between-deme transmission events over time.

### Predictors of H5N8 virus spread between poultry farms across borders

Alongside inferring the transmission dynamics of H5N8, potential drivers of virus spread between poultry farms across the four countries were investigated by quantifying the corresponding *R*_e_ parameter in the MTBD model with a generalized linear model (GLM) (26, 27) (see Materials and Methods). There were eight predictors included in the model: the 2016 live poultry trade (28), the 2016 poultry density in the source and destination deme (28), the 2014 poultry farm density in the source and destination deme (24), the 2017 farm outbreak density in the source and destination deme (25) and the distance between countries’ centroids (SI Appendix Table S5). The highest number of poultry was moved from Czech Republic (22 million) and Germany (20 million) to Poland. The highest poultry density was reported in Poland (600 birds/km^2^) while the highest poultry farm density was found in Germany (0.2 farm/km^2^). The highest poultry farm outbreak density was reported in Hungary (0.003 outbreak/km^2^). Countries centroids were 382 to 790 km apart. Figure 6A shows, for each predictor, the inclusion probability which represents the proportion of the posterior samples in which the given predictor was included in the model and the Bayes Factor (BF) which quantifies which of the posterior and prior inclusion probabilities of the given predictor in the model is more likely. Figure 6B shows the log conditional effect size which represents the log contribution of the given predictor when it was included in the model. None of the predictors were statistically supported to be associated with the spread of H5N8 virus between poultry farms across borders, illustrated by the low BF metric (<3.2) (Figure 6A) and the similar distribution between the posterior coefficient estimates (Figure 6B) and the prior (SI Appendix Table S2).

**Figure 6.**
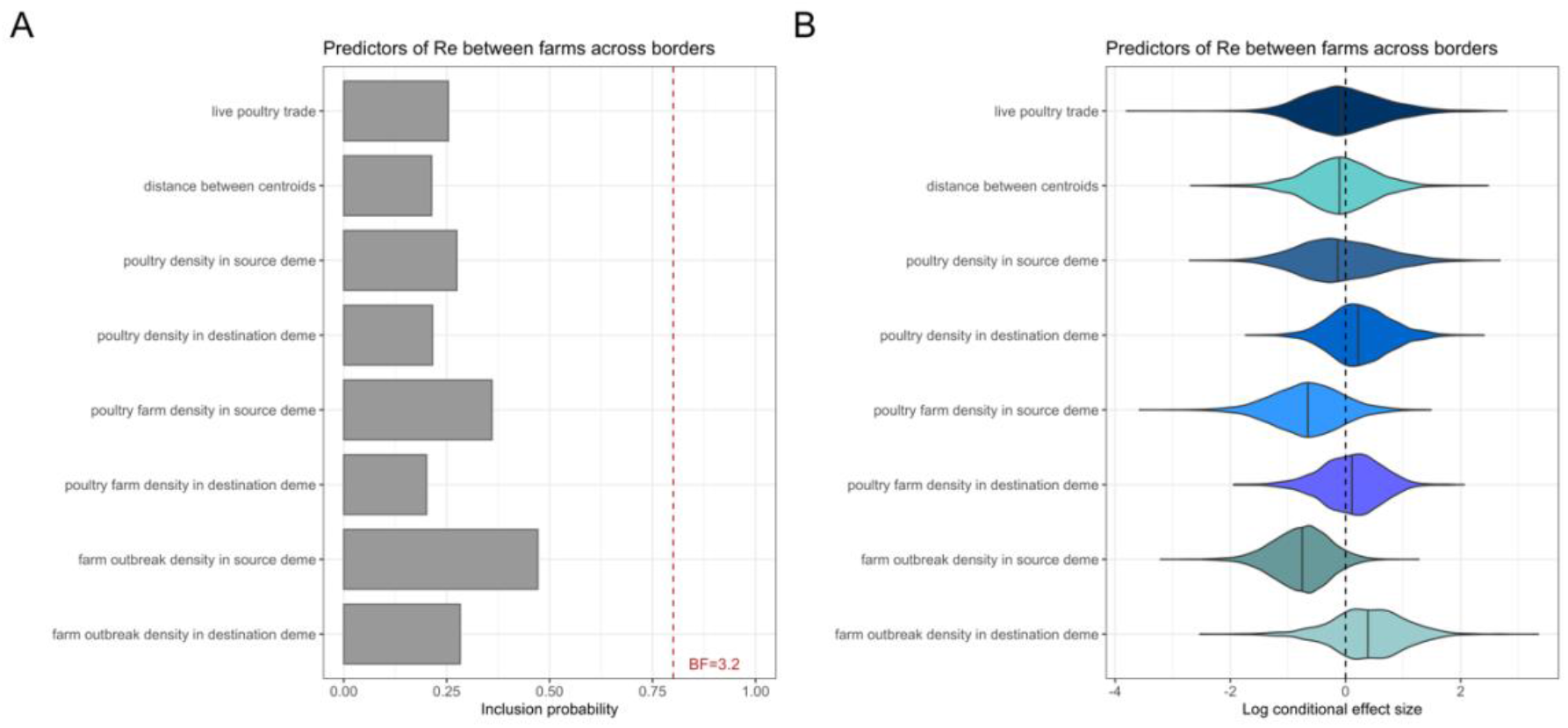
A) Inclusion probability for predictors of the between-farm H5N8 virus spread across borders. This represents the proportion of the posterior samples in which each predictor was included in the model. Bayes Factors (BF) were used to determine the contribution of each predictor in the GLM. BF were calculated for each predictor to quantify which of the posterior and prior inclusion probabilities of the given predictor in the model is more likely. The cutoff for substantial contribution of a given predictor in the GLM was set at 3.2 (51). B) Log conditional effect sizes for predictors of the between-farm H5N8 virus spread across borders. This represents the (log) contribution of each predictor when the corresponding predictor was included in the model (*β*_i_| *δ*_I_ = 1), where *β*_i_ is the coefficient and the binary indicator *δ*_I_ for each predictor *i*.

## Discussion

The epidemic trajectories showed that, in each country, the first introduction of H5N8 virus from wild birds to poultry farms likely occurred during autumn, which is in line with the timing of arrival of migratory wild birds in Europe (29). Also, the epidemic trajectories indicated that there was a delay of 3 to 16 weeks (depending of the country) between the inferred date of the first virus introduction and the date of the first officially reported poultry farm outbreak, likely illustrating different surveillance strategies’ effectiveness. The longest delay (16 weeks) was observed in Czech Republic, where most outbreaks occurred in small size farms (< 100 birds), while they mainly affected large size farms (> 10,000 birds) in Germany, Hungary and Poland (1). While a total of 442 poultry farm outbreaks and 372 wild bird cases in the four countries were officially reported (25), the epidemic trajectories showed that these numbers could have been under-estimated, especially in the wild bird population, likely due to challenges related to wildlife surveillance (30). High reporting rates of poultry farm outbreaks were found in Germany, Hungary and Poland, likely linked to the high mortality rates of poultry following H5N8 virus infection, along with the active surveillance implemented around reported poultry farm outbreaks (24). However, lower and again more delayed reporting rates were found for the poultry farms in Czech Republic. These results suggest that the likelihood of reporting infected farms is likely associated with the characteristics of the farm. However, whether this results of differences in size or other factors linked to the farm size (such as different farmers’ knowledge, attitudes and practices) needs further investigation.

Following the first virus introduction, the epidemic trajectories demonstrated that in Germany, Hungary and Poland, the epidemic was dominated by local farm-to-farm transmission events. In Germany, local farm-to-farm transmission increased between February and May 2017, likely illustrating the cluster of turkey farm outbreaks which represented approximately 25% of the total number of poultry farm outbreaks in the country (24). In Hungary, a peak of in the number of farm-to-farm transmission events was reported between October and November 2016, during which most outbreaks clustered in time and space (1). Moreover, the epidemic in these countries was also partly driven by wild bird-to-farm transmission (in particular in the middle of the epidemic) showing that the role of wild birds was likely greater than expected and was not limited to the onset of the epidemic. These outcomes also emphasize that in-place biosecurity measures in Germany, Hungary and Poland were sufficient to prevent continued incursions from farms across borders (such as ban of international trade) (24), but were less effective against local farm-to-farm and wild bird-to-farm transmission. Having more detailed knowledge of how poultry farms are connected with one another in those countries could help containing future outbreaks by disrupting the network of potential transmissions between poultry farms. Important efforts are also necessary to ensure that prevention strategies aiming at limiting the virus spread between wild birds and poultry (such as restriction of outdoor access and providing indoor feed and drinking water) (30) are implemented during high-risk periods. Also, more sustainable strategies should be explored for poultry farms for which access to outdoor areas is part of the production specifications. The contribution of wild birds to poultry farms outbreaks was even more substantial in Czech Republic, in which the epidemic trajectories showed that the epidemic was dominated by wild bird-to-farm transmission events. Accordingly, they also showed that the majority of farm-to-wild bird transmission events were from Czech Republic. This provides evidence that small size farms could be more exposed to virus transmission from wild birds than large commercial farms. Again, this could be explained by differences in farm size or other factors linked to the farm size (such as different farming practices – access to outdoor area – or biosecurity levels) which requires further attention. In wild birds, the epidemic was dominated by wild bird-to-wild bird transmission events. The number of wild bird-to-wild bird transmission events however decreased drastically from February 2017, likely linked to the decrease in wild bird density with migration to warmer climates (31) and the decrease in virus survival in the environment due to temperature-dependence of H5N8 virus transmission (24).

We also attempted to uncover factors that could potentially predict the spread of H5N8 virus between farms across countries. However, none of the investigated predictors were identified as supportive predictor of the viral spread. This is in line with outbreak investigations on affected poultry farms in Europe, which showed that the likelihood of H5N8 virus introduction from one country to another via personnel contacts, trade of live poultry, feed, or poultry products was negligible (7), although unreported cross-border activities could not be excluded. Using phylodynamic approaches, one previous study has found geographic proximity, sharing borders and live poultry trade (when using time-dependent predictors) to be strong drivers of AI virus spread between countries in Asia (32). The comparison of this previous GLM results to our study may not be appropriate due to differences in farming systems between Europe and Asia. Also, our predictors ignore other potential drivers of virus spread, such as wild bird migration, different farming systems and biosecurity levels among countries. For example, the scattered distribution of H5N8 sequences from wild birds among sequences from poultry farms of different countries on the MCC tree could support the possibility of wild birds’ movements facilitating virus spread between poultry farms across countries. It is also possible that transmission between countries is linked to trade of poultry products or other cryptic means that were not tested in this study due to lack of information. In the future, we recommend further investigation of predictors with a higher scale of temporal and spatial resolutions, which could allow for stronger contribution levels (32).

The 2016-2017 epidemic of H5N8 virus in Europe remains, like other epidemics of AI viruses, epidemiologically complex as it involved multiple wild bird species that vary in spatial ecology and clinical disease severity. During the epidemic, the virus was detected in a large number of wild bird species, mainly those of the *Anseriformes* orders (ducks, geese, swans), including mute swans (*Cygnus olor*), tufted ducks (*Aythya fuligula*), Whooper swans (*Cygnus cygnus*), Eurasian widgeons (*Mareca penelope*) and mallards (*Anas platyrhynchos*) (24). Among these species, some can be mostly sedentary in given areas while partially or wholly migratory in others (29), meaning that some species can act as sentinels in some areas or long-distance vectors of H5N8 virus in others (33). Consequently, wild bird population structure may be much more complex than what was assumed in this study. For example, on the MCC tree, we observed H5N8 sequences from wild birds both within and between clusters of sequences from poultry farms of the same country, which could illustrate the presence of sedentary and migratory wild birds, respectively. Similarly, the virus was detected in several poultry species and farm types, which may play different roles in the virus spread due to discrepancies in virus infection susceptibility and farming practices (1, 24). Unfortunately, limited information on virus prevalence or epidemiology in various domestic and wild host species between countries makes it difficult to treat species separately, thereby necessitating the grouping used here.

Bayesian phylogeographic approaches (34) are more common than structured phylodynamic approaches (like the MTBD model) to infer the transmission of lineages between different host species, their popularity being partly associated to their computational efficiency (7, 35). However, one shortcoming of Bayesian phylogeographic approaches is the assumption of independence between the phylogeny and the transmission process, which can lead to loss of information (36–38). Another shortcoming is the assumption of proportionality between the sample sizes across sub-populations and the subpopulation sizes, which make it sensitive to biased sampling. Unlike Bayesian phylogeographic approaches, MTBD models explicitly integrate how lineages transmit within and between sub-populations while accounting for the sampling effort, making the estimations more robust to sampling bias (19, 21). This has made it possible to infer transmission parameters, such as the effective reproduction number *R*_e_, among poultry farms and wild birds based on pathogen genome sequences. We expect those estimates being useful to parameterize predictive models of virus spread aiming at optimising control strategies. We also inferred that the median farm-level infectious periods ranged from 7 to 14 days, suggesting that some countries were quicker at depopulating than others. This also emphasizes that a back-tracing window of approximately 2 weeks would be sufficient to capture the period during which a farm was infectious. Only one sequence was available per poultry farm, meaning that within-farm genetic diversity was not taken into account. However, this is a reasonable assumption due to the short-period of the poultry farm outbreaks prior to detection and culling. More importantly, the present study demonstrates how relevant these models can be (i) to inform on the number of unreported infections, (ii) to reconstruct previous unobserved infections prior to the first officially reported infection and (iii) to discriminate transmission events within a given host species from incursions across species, that are more challenging using traditional wildlife and epidemiological methods (15, 39). Therefore, such phylodynamic tools can complement or even substitute for traditional epidemiological tools.

Phylodynamics provides one avenue for quantifying patterns and identifying drivers of infectious disease transmission dynamics at the wildlife-domestic animal interface, which is a fundamental challenge for veterinary epidemiology. We expect our results will be valuable in better informing policy decision-making as means to reduce the impact of future epidemics of HPAI viruses.

## Methods

### Selection and alignment of sequences

All H5N8 genome sequences of HA segment collected during winter 2016-2017 from four severely affected European countries (Czech Republic, Germany, Hungary and Poland) were downloaded from GISAID on Sept 1^st^, 2020. Only one sequence was available per poultry farm and per wild bird, meaning that the transmission dynamics of H5N8 were inferred at the farm-to-farm, wild bird-to-farm and farm-to-wild bird levels. Selected sequences were annotated with available sampling dates, locations and hosts, aligned using MAFFT v7 (40) and manually edited using AliView v1.26 (41). The final dataset consisted of 190 genome sequences from infected poultry farms and 130 from infected wild birds (SI Appendix Table S6 and Figure S3).

### Phylodynamic analysis

#### Multi-type birth-death model

The multi-type birth-death (MTBD) model was fitted to the sequence alignment (19, 21). Under this model, infected hosts could transmit the virus to another host from the same discrete subpopulation (with a parameter within-deme *R*_e_), referred to here as deme, eventually become non-infectious due to recovery or death/depopulation (with a rate δ), be sequenced and sampled upon becoming non-infectious (with a proportion *s*, and thus are included into the dataset) or could transmit the virus to another host from another deme (with a parameter between-deme *R*_e_). All transmission, become non-infectious and sampling processes are assumed to be deme-specific and constant through time, except for the within-deme *R*_e_ that was estimated across four-time intervals, corresponding to the four phases of the epidemic (SI Appendix Figure S1).

Under this model, sequences were organized into five demes, according to the host type and geographical location: ‘poultry farms in Czech Republic’, ‘poultry farms in Germany’, ‘poultry farms in Hungary’, ‘poultry farms in Poland’ and ‘wild birds in the four countries’ (SI Appendix Table S6). All sequences from wild birds were aggregated into one deme (not depending on the geographical location as for poultry farms) since it was assumed that the majority of sampled wild bird species (mainly mallards and swans) could move freely among countries (42). It was assumed that once sampled, a given host could not be infected and sampled again since infected poultry farms were subject to culling following the confirmation of infection and sampling of wild birds was from a mortality event (24).

The prior values and distributions of the model parameters are described in SI Appendix Table S2. The MTBD was specified as the tree prior using a HKY + Γ4 nucleotide substitution process with a relaxed molecular clock (43) defined by a Lognormal(0.001, 1.25) prior (4, 5, 44). The origin of the tree was given a Lognormal(−0.2,0.2) prior, corresponding to the median date 1^st^ of July 2016 (95% HPD: 6^th^ of February 2016 – 19^th^ of October 2016) (2, 4, 5) and assumed to be associated to the deme ‘wild birds in the four countries’, since the source of the first poultry farm outbreak in the four countries was likely attributed to infected migratory wild birds from Northern Eurasia (24). All *R*_e_ parameters were given a Lognormal(0,1) prior (9, 45, 46). The become non-infectious rate was given a Lognormal(52,0.6) prior (46–48). For each deme, the sampling proportion was given a Uniform distribution prior with lower and upper bounds informed by the number of sequences and reported poultry farm outbreaks/wild bird cases (25). Given the severity of the clinical signs affecting the majority of poultry combined with active surveillance around reported poultry farm outbreaks (24), the number of unreported poultry farm outbreaks was considered relatively low in all countries. On the contrary, given the difficulty of catching and sampling wild birds, it was assumed that infected wild birds were significantly under-sampled, relative to poultry farms.

#### Predictors of H5N8 virus spread between poultry farms across borders

The MTBD model was extended with a generalized linear model (GLM) to inform the H5N8 virus spread between poultry farms across borders by 19 time-independent predictors (26, 27): the 2016 live poultry trade (28), the 2016 poultry density in the source and destination deme (28), the 2014 poultry farm density in the source and destination deme (24), the 2017 farm outbreak density in the source and destination deme (25), the 2021 human density in the source and destination deme (49), whether two countries shared borders and the distance between countries’ centroids. To account for potential missing predictors, we also included predictors to assess the virus spread from or to one individual country (SI Appendix Table S5). In this GLM parametrization, the between-deme *R*_e_ parameters act as the outcome to a log-linear function of the predictors. For each predictor *i*, the GLM parametrization also includes a regression coefficient *β*_i_ which quantifies the (log) contribution of the predictor and a binary indicator variable *δ*_i_ which quantifies the probability of the predictor to be included in the model (SI Appendix Table S2). To avoid collinearity among predictors, predictors were removed when the Pearson correlation exceeded > 0.7 (SI Appendix Figure S4). To reduce the effect of different predictors’ magnitude, all non-binary predictors were log-transformed and standardized before inclusion in the GLM. Bayes Factors (BF) were used to determine the contribution of each predictor in the GLM (26, 50, 51). BF were calculated for each predictor to quantify which of the posterior and prior inclusion probabilities of the given predictor in the model (*δ*_I_ = 1) is more likely. The cutoff for substantial contribution of a given predictor in the GLM was set at 3.2 (51), meaning that its posterior inclusion probability in the model was 3.2-fold more likely than its prior inclusion probability (0.50).

#### Inference of MTBD model parameters, structured trees and epidemic trajectories

Phylodynamic analysis was implemented using the BDMM-Prime package (52) for BEAST v2.6.3 (53) and the BEAGLE library (54) to improve computational performance. All analyses were run for 40-50 million steps across three independent Markov chains (MCMC) and states were sampled every 10,000 steps. The first 10% of steps from each analysis were discarded as burn-in before states from the chains were pooled using Log-Combiner v2.6.3 (53). Convergence was assessed in Tracer v1.7 (55) by ensuring that the estimated sampling size (ESS) values associated with the estimated parameters were all > 200.

The structured trees (i.e. when phylogenetic trees are associated with a specific deme along their branches (22)) were inferred by applying a stochastic mapping algorithm (56) implemented in BDMM-Prime (52) to a subsampled set of posterior structured trees and model parameters (n=500) generated by the MTBD analysis. The MCC tree was obtained from the structured trees in TreeAnnotator v2.6.3 (53), and annotated using the ggtree package (57) in R v4.0.2 (58). For each set of posterior model parameters set and associated structured tree, an epidemic trajectory (i.e. corresponding to the sequence of transmission, become non-infectious and sampling events that occur throughout a given epidemic (23)) was drawn from the distribution of such trajectories conditional on the tree and parameters (52).

## Data availability

All H5N8 genome sequences of HA segment are available on the GISAID database (https://www.gisaid.org). The prior values and distributions of the model parameters are described in SI Appendix Table S2. Details on the predictor data are available in SI Appendix Table S5. The BEAST 2 XML file used to perform the phylodynamic analysis, together with the R scripts are available from https://github.com/ClaireGuinat/h5n8_bdmm-prime.git.

## Acknowledgements

The authors are grateful to the cEvo group (ETH Zurich, Switzerland) for providing useful comments on this project. They also acknowledge the EFSA Public Access to documents Team for providing data on the poultry farm census in Europe. This project has received funding from the European Union’s Horizon 2020 research and innovation programme under the Marie Sklodowska-Curie grant agreement No 842621.

## Competing interests

The authors declare no competing interests.

## Notes

### Competing Interest Statement

The authors have declared no competing interest.

## References

1. S. Napp, N. Majó, R. Sánchez-Gónzalez, J. Vergara-Alert, Emergence and spread of highly pathogenic avian influenza A(H5N8) in Europe in 2016-2017. Transbound. Emerg. Dis. 65, 1217–1226 (2018).

2. E. Świętoń, K. Śmietanka, Phylogenetic and molecular analysis of highly pathogenic avian influenza H5N8 and H5N5 viruses detected in Poland in 2016–2017. Transbound. Emerg. Dis. 65, 1664–1670 (2018).

3. A. Globig, et al., Highly Pathogenic Avian Influenza H5N8 Clade 2.3.4.4b in Germany in 2016/2017. Front. Vet. Sci. 4 (2018).

4. A. Fusaro, et al., Genetic Diversity of Highly Pathogenic Avian Influenza A(H5N8/H5N5) Viruses in Italy, 2016–17. Emerg. Infect. Dis. 23, 1543–1547 (2017).

5. N. Beerens, et al., Multiple Reassorted Viruses as Cause of Highly Pathogenic Avian Influenza A(H5N8) Virus Epidemic, the Netherlands, 2016. Emerg. Infect. Dis. 23, 1974–1981 (2017).

6. P. Mulatti, et al., Integration of genetic and epidemiological data to infer H5N8 HPAI virus transmission dynamics during the 2016-2017 epidemic in Italy. Sci. Rep. 8, 18037 (2018).

7. S. Lycett, et al., Global Consortium for H5N8 and Related Influenza Viruses. Role Migr. Wild Birds Glob. Spread Avian Influenza H5N8 Sci. 354, 213–217 (2016).

8. C. Guinat, et al., Role of Live-Duck Movement Networks in Transmission of Avian Influenza, France, 2016–2017. Emerg. Infect. Dis. 26, 472–480 (2020).

9. A. Andronico, et al., Highly pathogenic avian influenza H5N8 in south-west France 2016–2017: A modeling study of control strategies. Epidemics 28, 100340 (2019).

10. C. Guinat, et al., Biosecurity risk factors for highly pathogenic avian influenza (H5N8) virus infection in duck farms, France. Transbound. Emerg. Dis. 67, 2961–2970 (2020).

11. C. Guinat, N. Rouchy, F. Camy, J. L. Guérin, M. C. Paul, Exploring the Wind-Borne Spread of Highly Pathogenic Avian Influenza H5N8 During the 2016–2017 Epizootic in France. Avian Dis. 63, 246–248 (2018).

12. A. Scoizec, et al., Airborne Detection of H5N8 Highly Pathogenic Avian Influenza Virus Genome in Poultry Farms, France. Front. Vet. Sci. 5 (2018).

13. E. M. Volz, K. Koelle, T. Bedford, Viral phylodynamics. PLoS Comput Biol 9, e1002947 (2013).

14. L. du Plessis, T. Stadler, Getting to the root of epidemic spread with phylodynamic analysis of genomic data. Trends Microbiol. 23, 383–386 (2015).

15. G. Guinat, et al., What can phylodynamics bring to animal health research? Trends Ecol. Evol. (2021).

16. G. Dudas, L. M. Carvalho, A. Rambaut, T. Bedford, MERS-CoV spillover at the camel-human interface. Elife 7, e31257 (2018).

17. N. R. Faria, et al., Genomic and epidemiological monitoring of yellow fever virus transmission potential. Science 361, 894–899 (2018).

18. S. A. Nadeau, T. G. Vaughan, J. Scire, J. S. Huisman, T. Stadler, The origin and early spread of SARS-CoV-2 in Europe. Proc. Natl. Acad. Sci. 118 (2021).

19. D. Kühnert, T. Stadler, T. G. Vaughan, A. J. Drummond, Phylodynamics with migration: a computational framework to quantify population structure from genomic data. Mol. Biol. Evol. 33, 2102–2116 (2016).

20. R. M. Anderson, R. M. May, Population biology of infectious diseases: Part I. Nature 280, 361–367 (1979).

21. J. Scire, J. Barido-Sottani, D. Kühnert, T. G. Vaughan, T. Stadler, Improved multi-type birth-death phylodynamic inference in BEAST 2. BioRxiv (2020).

22. T. G. Vaughan, D. Kühnert, A. Popinga, D. Welch, A. J. Drummond, Efficient Bayesian inference under the structured coalescent. Bioinforma. Oxf. Engl. 30, 2272–2279 (2014).

23. T. G. Vaughan, et al., Estimating epidemic incidence and prevalence from genomic data. Mol. Biol. Evol. 36, 1804–1816 (2019).

24. EFSA, et al., Avian influenza overview October 2016–August 2017. EFSA J. 15, e05018 (2017).

25. FAO, Global Animal Disease Information System (Empres-i) [Available at: https://empres-i.review.fao.org/#/ (accessed in January 2021)] (2021).

26. P. Lemey, et al., Unifying viral genetics and human transportation data to predict the global transmission dynamics of human influenza H3N2. PloS Pathog 10, e1003932 (2014).

27. N. F. Müller, G. Dudas, T. Stadler, Inferring time-dependent migration and coalescence patterns from genetic sequence and predictor data in structured populations. Virus Evol. 5, vez030 (2019).

28. FAOSTAT, FAOSTAT - production and trade of live animals [Available at: http://www.fao.org/faostat/en/#compare (accessed in May 2021)] (2021).

29. BTO, British Trust for Ornithology (BTO) <https://www.bto.org> [accessed in May 2021] (2017).

30. M. Artois, et al., Outbreaks of highly pathogenic avian influenza in Europe: the risks associated with wild birds. Rev. Sci. Tech. 28, 69 (2009).

31. N. J. Hill, et al., Transmission of influenza reflects seasonality of wild birds across the annual cycle. Ecol. Lett. 19, 915–925 (2016).

32. J. Yang, D. Xie, Z. Nie, B. Xu, A. J. Drummond, Inferring host roles in bayesian phylodynamics of global avian influenza A virus H9N2. Virology 538, 86–96 (2019).

33. J. Keawcharoen, et al., Wild ducks as long-distance vectors of highly pathogenic avian influenza virus (H5N1). Emerg. Infect. Dis. 14, 600 (2008).

34. P. Lemey, A. Rambaut, A. J. Drummond, M. A. Suchard, Bayesian Phylogeography Finds Its Roots. PLOS Comput. Biol. 5, e1000520 (2009).

35. N. S. Trovão, M. A. Suchard, G. Baele, M. Gilbert, P. Lemey, Bayesian Inference Reveals Host-Specific Contributions to the Epidemic Expansion of Influenza A H5N1. Mol. Biol. Evol. 32, 3264–3275 (2015).

36. N. De Maio, C.-H. Wu, K. M. O’Reilly, D. Wilson, New Routes to Phylogeography: A Bayesian Structured Coalescent Approximation. PLoS Genet. 11, e1005421 (2015).

37. S. Bloomfield, et al., Investigation of the validity of two Bayesian ancestral state reconstruction models for estimating Salmonella transmission during outbreaks. Plos One 14, e0214169 (2019).

38. N. F. Müller, D. A. Rasmussen, T. Stadler, The Structured Coalescent and Its Approximations. Mol. Biol. Evol. 34, 2970–2981 (2017).

39. J. A. Blanchong, S. J. Robinson, M. D. Samuel, J. T. Foster, Application of genetics and genomics to wildlife epidemiology. J. Wildl. Manag. 80, 593–608 (2016).

40. K. Katoh, D. M. Standley, MAFFT Multiple Sequence Alignment Software Version 7: Improvements in Performance and Usability. Mol. Biol. Evol. 30, 772–780 (2013).

41. Larsson, AliView: a fast and lightweight alignment viewer and editor for large datasets. Bioinformatics 30, 3276–3278 (2014).

42. P. W. Atkinson, et al., “Urgent preliminary assessment of ornithological data relevant to the spread of Avian Influenze in Europe” (Wetlands International, 2006).

43. A. J. Drummond, S. Y. W. Ho, M. J. Phillips, A. Rambaut, Relaxed phylogenetics and dating with confidence. PLoS Biol. 4, e88 (2006).

44. P. Alarcon, et al., Comparison of 2016–17 and previous epizootics of highly pathogenic avian influenza H5 Guangdong lineage in Europe. Emerg. Infect. Dis. 24, 2270 (2018).

45. I. Iglesias, et al., Reproductive ratio for the local spread of highly pathogenic avian influenza in wild bird populations of Europe, 2005–2008. Epidemiol. Infect. 139, 99–104 (2011).

46. D. A. Grear, J. S. Hall, R. J. Dusek, H. S. Ip, Inferring epidemiologic dynamics from viral evolution: 2014-2015 Eurasian/North American highly pathogenic avian influenza viruses exceed transmission threshold, R0 = 1, in wild birds and poultry in North America. Evol. Appl. 11, 547–557 (2018).

47. C. Leyson, et al., Pathogenicity and genomic changes of a 2016 European H5N8 highly pathogenic avian influenza virus (clade 2.3. 4.4) in experimentally infected mallards and chickens. Virology 537, 172–185 (2019).

48. K. Willgert, et al., Transmission of highly pathogenic avian influenza in the nomadic free-grazing duck production system in Viet Nam. Sci. Rep. 10, 1–11 (2020).

49. Wikipedia, World population [Available at: https://en.wikipedia.org/wiki/World_population (accessed in July 2021)] (2021).

50. D. Magee, R. Beard, M. A. Suchard, P. Lemey, M. Scotch, Combining phylogeography and spatial epidemiology to uncover predictors of H5N1 influenza A virus diffusion. Arch. Virol. 160, 215–224 (2015).

51. R. E. Kass, A. E. Raftery, Bayes Factors. J. Am. Stat. Assoc. 90, 773–795 (1995).

52. T. Vaughan, BDMM Prime (In prep.).

53. R. Bouckaert, et al., BEAST 2: a software platform for Bayesian evolutionary analysis. PLoS Comput Biol 10, e1003537 (2014).

54. D. L. Ayres, et al., BEAGLE: an application programming interface and high-performance computing library for statistical phylogenetics. Syst. Biol. 61, 170–173 (2012).

55. A. Rambaut, A. J. Drummond, D. Xie, G. Baele, M. A. Suchard, Posterior summarization in Bayesian phylogenetics using Tracer 1.7. Syst. Biol. 67, 901 (2018).

56. W. A. Freyman, S. Höhna, Stochastic character mapping of state-dependent diversification reveals the tempo of evolutionary decline in self-compatible Onagraceae lineages. Syst. Biol. 68, 505–519 (2019).

57. G. Yu, D. K. Smith, H. Zhu, Y. Guan, T. T. Lam, ggtree: an R package for visualization and annotation of phylogenetic trees with their covariates and other associated data. Methods Ecol. Evol. 8, 28–36 (2017).

58. R. C. Team, R: A language and environment for statistical computing (2013).

