## Supplementary files for "Disentangling the role of poultry farms and wild birds in the spread of highly pathogenic avian influenza virus H5N8 in Europe"

### Supplementary Information

### Tables

Table S1. First transmission events extracted from the epidemic trajectories

| Deme | Date of the first officially reported outbreak/case | Inferred median date of the first imported (i.e. from another deme) outbreak/case (95% High Posterior Density) | Median time delay between the date of the first officially reported outbreak/case and the inferred date of the first imported outbreak/case (days) (95% High Posterior Density) | Inferred median date of the first local (i.e. within-deme) outbreak/case (95% High Posterior Density) | Median time delay between the date of the first officially reported outbreak/case and the inferred date of the first local outbreak/case (days) (95% High Posterior Density) | Inferred median date of the first exported (i.e. to another deme) outbreak/case (95% High Posterior Density) | Median time delay between the date of the first officially reported outbreak/case and the inferred date of the first exported outbreak/case (days) (95% High Posterior Density) |
|---|---|---|---|---|---|---|---|
| Poultry farms in Czech Republic | 5/01/2017 | 11/09/2016 (30/07/2016 – 27/10/2016) | 115 (70 – 158) | 20/09/2016 (10/08/2016 – 08/12/2016) | 101 (26 – 146) | 12/09/2016 (01/08/2016 – 01/11/2016) | 114 (63 – 155) |
| Poultry farms in Germany | 10/11/2016 | 20/10/2016 (17/09/2016 – 05/11/2016) | 21 (5 – 54) | 29/10/2016 (08/10/2016 – 07/11/2016) | 12 (-1 – 32) | 28/10/2016 (06/10/2016- 14/11/2016) | 13 (-5 – 35) |
| Poultry farms in Hungary | 4/11/2016 | 01/10/2016 (17/08/2016 – 18/10/2016) | 34 (14 – 77) | 08/10/2016 (31/08/2016 – 25/10/2016) | 29 (9 – 65) | 14/10/2016 (04/09/2016 – 10/11/2016) | 20.5 (-7 – 59) |
| Poultry farms in Poland | 5/12/2016 | 20/10/2016 (20/08/2016 – 29/11/2016) | 46 (5 – 106) | 11/11/2016 (29/08/2016 – 05/12/2016) | 24 (1 – 97) | 03/11/2016 (29/08/2016 – 16/12/2016) | 31.5 (-11 – 97) |
| Wild birds in the four countries | 27/10/2016 | 09/09/2016 (01/08/2016 – 16/10/2016) | 48 (10 – 86) | 17/08/2016 (02/07/2016 – 19/08/2016) | 71 (38 – 116) | 04/09/2016 (28/07/2016 – 08/10/2016) | 53 (15 – 87) |

Table S2. Prior values and distributions used in the multi-type birth-death model

| Parameter | Prior | Rationale | References |
|---|---|---|---|
| Nucleotide substitution model | HKY + $\Gamma_4$ | Unequal transition/transversion rates, unequal base frequencies, rate heterogeneity among sites with four categories | (1–3) |
| Clock rate | Lognormal(0.001, 1.25) | Median of 0.0004 substitution.site$^{-1}$.year$^{-1}$ [95% interquartile range (IQR): 3.10$^{-5}$-0.005] | (1, 3, 3) |
| Effective reproduction number within each deme | Lognormal(0,1) | Median of 1 [95%IQR: 0.1-7.1] | (4–6) |
| Effective reproduction | Lognormal(0,1) | Median of 1 [95%IQR: 0.1-7.1] | (4–6) |

| | | | |
|---|---|---|---|
| number between demes | | | |
| Become uninfectious rate | Lognormal(52,0.6) | Median of 43.4 $year^{-1}$ [95% IQR: 13.4-141] | (4, 7, 8) |
| Sampling proportion | | Upper bounds are defined by the number of sequences over the number of reported outbreaks/cases; lower bounds are defined by the number of sequences over the number of reported outbreaks/cases*1.5 | (4, 7, 8) |
| Poultry farms in Czech Republic | Uniform(0.553,0.837) | 36/65 - 36/43 | (9) |
| Poultry farms in Germany | Uniform(0.560,0.840) | 79/141 - 79/94 | (9) |
| Poultry farms in Hungary | Uniform(0.083,0.125) | 30/360 - 30/240 | (9) |
| Poultry farms in Poland | Uniform(0.459,0.692) | 45/98 - 45/65 | (9) |
| Wild birds in the four countries | Uniform(0,0.349) | 0 - 130/372 | (9) |
| Time of origin | Lognormal(-0.2,0.2) | Median of 1 Juillet 2016 [95% HPD: 6 February 2016 - 19 October 2016] | (2, 3, 10) |
| Deme of origin | Wild birds in the four countries | Putative epidemic origin | (11) |
| Regression coefficient | Normal(0,2) | | (12) |
| Indicator variable | 0 or 1 | 0.5 prior inclusion probability for each predictor | (12) |

Table S3. Posterior values and distributions of key epidemiological parameters inferred by the multi-type birth-death model

| Parameter | Posterior median | 95% High Posterior Density |
|---|---|---|
| **Within-deme effective reproduction number** | | |
| **Poultry farms in Czech Republic** | | |
| Period 1 (Oct-Nov 2016) | 0.8 | 0.0 - 3.5 |
| Period 2 (Dec 2016) | 1.1 | 0.1 - 2.8 |
| Period 3 (Jan 2017) | 0.3 | 0.0 - 0.7 |
| Period 4 (Feb-May 2017) | 0.2 | 0.0 - 0.5 |
| **Poultry farms in Germany** | | |
| Period 1 (Oct-Nov 2016) | 0.9 | 0.4 - 1.7 |
| Period 2 (Dec 2016) | 0.7 | 0.3 – 1.2 |
| Period 3 (Jan 2017) | 0.5 | 0.2 – 0.8 |

| | | |
|---|---|---|
| Period 4 (Feb-May 2017) | 0.9 | 0.6 – 1.3 |
| **Poultry farms in Hungary** | | |
| Period 1 (Oct-Nov 2016) | 1.3 | 0.8 – 1.8 |
| Period 2 (Dec 2016) | 0.5 | 0.1 – 1.0 |
| Period 3 (Jan 2017) | 0.4 | 0.1 – 0.8 |
| Period 4 (Feb-May 2017) | 0.6 | 0.3 – 1.1 |
| **Poultry farms in Poland** | | |
| Period 1 (Oct-Nov 2016) | 0.8 | 0.1 – 2.7 |
| Period 2 (Dec 2016) | 0.9 | 0.5 – 1.3 |
| Period 3 (Jan 2017) | 0.3 | 0.1 – 0.6 |
| Period 4 (Feb-May 2017) | 0.5 | 0.2 – 1.0 |
| **Wild birds in the four countries** | | |
| Period 1 (Oct-Nov 2016) | 1.4 | 1.1 – 1.7 |
| Period 2 (Dec 2016) | 1.3 | 0.9 – 1.7 |
| Period 3 (Jan 2017) | 1.1 | 0.8 – 1.4 |
| Period 4 (Feb-May 2017) | 0.1 | 0.0 – 0.2 |
| **Between-deme effective reproduction number** | | |
| From poultry farms in Czech Republic | | |
| To poultry farms in Germany | 6.9.10-3 | 2.3.10-2 – 2.2.10-5 |
| To poultry farms in Hungary | 4.6.10-3 | 1.4.10-2 – 1.6.10-5 |
| To poultry farms in Poland | 1.1.10-2 | 4.4.10-2 – 9.3.10-5 |
| To wild birds in the four countries | 1.4.10-2 | 2.8.10-2 – 5.2.10-3 |
| From poultry farms in Germany | | |
| To poultry farms in Czech Republic | 9.7.10-3 | 4.3.10-2 – 7.3.10-5 |
| To poultry farms in Hungary | 4.5.10-3 | 1.4.10-2 – 1.3.10-5 |
| To poultry farms in Poland | 1.1.10-2 | 5.2.10-2 – 4.4.10-5 |
| To wild birds in the four countries | 1.3.10-2 | 2.6.10-2 – 5.6.10-3 |
| From poultry farms in Hungary | | |
| To poultry farms in Czech Republic | 1.1.10-2 | 7.5.10-2 – 2.2.10-5 |
| To poultry farms in Germany | 8.2.10-3 | 3.9.10-2 – 1.1.10-6 |
| To poultry farms in Poland | 1.3.10-2 | 7.3.10-2 – 4.1.10-5 |
| To wild birds in the four countries | 2.4.10-2 | 5.0.10-2 – 7.6.10-3 |
| From poultry farms in Poland | | |
| To poultry farms in Czech Republic | 9.3.10-3 | 5.0.10-2 – 3.9.10-5 |
| To poultry farms in Germany | 6.8.10-3 | 2.5.10-2 – 2.4.10-5 |
| To poultry farms in Hungary | 4.5.10-3 | 1.4.10-2 – 2.6.10-5 |
| To wild birds in the four countries | 1.5.10-2 | 2.9.10-2 – 5.6.10-3 |
| From wild birds in the four countries | | |
| To poultry farms in Czech Republic | 4.6 | 0.9 – 9.2 |
| To poultry farms in Germany | 0.2 | 0.0 – 0.5 |
| To poultry farms in Hungary | 0.1 | 0.0 – 0.3 |
| To poultry farms in Poland | 0.4 | 0.0 – 1.3 |
| **Infectious period (days)** | | |
| Poultry farms in Czech Republic | 14 | 7 - 23 |
| Poultry farms in Germany | 10 | 6 - 16 |
| Poultry farms in Hungary | 8 | 4 - 14 |
| Poultry farms in Poland | 7 | 4 - 11 |
| Wild birds in the four countries | 14 | 11-19 |

Table S4. Median value of the cumulative number of outbreaks/cases due to local virus transmission (i.e. within-deme) versus importations (i.e. between-deme) (95% High Posterior Density) extracted from the epidemic trajectories

| Deme | Inferred median cumulative number of outbreaks/cases due to local virus transmission (i.e. within-deme) versus importations (i.e. between-deme) (95% High Posterior Density) | | | | |
|---|---|---|---|---|---|
| | Poultry farms in Czech Republic | Poultry farms in Germany | Poultry farms in Hungary | Poultry farms in Poland | Wild birds in the four countries |
| Poultry farms in Czech Republic | 56 (3 – 205) | 2 (1 – 80) | 2 (1 – 10) | 2 (1 – 18) | 115 (57 – 230) |
| Poultry farms in Germany | 2 (1 – 14) | 109 (61 – 5750) | 2 (1 – 8) | 3 (1 – 13) | 50 (22 – 136) |
| Poultry farms in Hungary | 4 (1 – 28) | 3 (1 – 60) | 316 (144 – 888) | 3 (1 – 29) | 101 (27 – 291) |
| Poultry farms in Poland | 3 (1 – 16) | 2 (1 – 86) | 2 (1 – 9) | 77 (32 – 367) | 59 (27 – 148) |
| Wild birds in the four countries | 972 (77 – 5,569) | 47 (7 – 1573) | 68 (11 – 207) | 77 (2 – 422) | 3,075 (807 – 8,575) |

Table S5. Information related to the predictors used in this study. CR: Czech Republic, GE: Germany, HU: Hungary, PO: Poland

| Trade of live poultry between countries | | | | |
|---|---|---|---|---|
| | CR | GE | HU | PO |
| CR | | 9239000 | 42000 | 21659000 |
| GE | 8745000 | | 1750000 | 20422000 |
| HU | 1255000 | 406000 | | 2093000 |
| PO | 509000 | 5802000 | 20000 | |
| **Shared border** | | | | |
| | CR | GE | HU | PO |
| CR | | 1 | 0 | 1 |
| GE | 1 | | 0 | 1 |
| HU | 0 | 0 | | 0 |
| PO | 1 | 1 | 0 | |
| **Distance between centroids (km)** | | | | |

| | CR | GE | HU | PO |
|---|---|---|---|---|
| CR | | 382.7534 | 415.9354 | 389.9038 |
| GE | 382.7534 | | 790.3776 | 633.6363 |
| HU | 415.9354 | 790.3776 | | 551.4372 |
| PO | 389.9038 | 633.6363 | 551.4372 | |

Density of human population in source deme

| | CR | GE | HU | PO |
|---|---|---|---|---|
| CR | | 136.03758 | 136.03758 | 136.03758 |
| GE | 234.435434 | | 234.435434 | 234.435434 |
| HU | 103.841245 | 103.841245 | | 103.841245 |
| PO | 121.039824 | 121.039824 | 121.039824 | |

Density of human population in destination deme

| | CR | GE | HU | PO |
|---|---|---|---|---|
| CR | | 234.435434 | 103.841245 | 121.039824 |
| GE | 136.03758 | | 103.841245 | 121.039824 |
| HU | 136.03758 | 234.435434 | | 121.039824 |
| PO | 136.03758 | 234.435434 | 103.841245 | |

Density of poultry in source deme

| | CR | GE | HU | PO |
|---|---|---|---|---|
| CR | | 270.251423 | 270.251423 | 270.251423 |
| GE | 488.9839 | | 488.9839 | 488.9839 |
| HU | 433.559067 | 433.559067 | | 433.559067 |
| PO | 599.985928 | 599.985928 | 599.985928 | |

Density of poultry in destination deme

| | CR | GE | HU | PO |
|---|---|---|---|---|
| CR | | 488.9839 | 433.559067 | 599.985928 |
| GE | 270.251423 | | 433.559067 | 599.985928 |
| HU | 270.251423 | 488.9839 | | 599.985928 |
| PO | 270.251423 | 488.9839 | 433.559067 | |

Density of poultry farms in source deme

| | CR | GE | HU | PO |
|---|---|---|---|---|
| CR | | 0.00426012 | 0.00426012 | 0.00426012 |
| GE | 0.21971482 | | 0.21971482 | 0.21971482 |
| HU | 0.02168118 | 0.02168118 | | 0.02168118 |
| PO | 0.00990153 | 0.00990153 | 0.00990153 | |

Density of poultry farms in destination deme

| | CR | GE | HU | PO |
|---|---|---|---|---|
| CR | | 0.21971482 | 0.02168118 | 0.00990153 |
| GE | 0.00426012 | | 0.02168118 | 0.00990153 |
| HU | 0.00426012 | 0.21971482 | | 0.00990153 |
| PO | 0.00426012 | 0.21971482 | 0.02168118 | |

| Density of poultry farm outbreaks in source deme | | | | |
|---|---|---|---|---|
| | CR | GE | HU | PO |
| CR | | 0.00054519 | 0.00054519 | 0.00054519 |
| GE | 0.00026302 | | 0.00026302 | 0.00026302 |
| HU | 0.00257981 | 0.00257981 | | 0.00257981 |
| PO | 0.00020788 | 0.00020788 | 0.00020788 | |

| Density of poultry farm outbreaks in destination deme | | | | |
|---|---|---|---|---|
| | CR | GE | HU | PO |
| CR | | 0.00026302 | 0.00257981 | 0.00020788 |
| GE | 0.00054519 | | 0.00257981 | 0.00020788 |
| HU | 0.00054519 | 0.00026302 | | 0.00020788 |
| PO | 0.00054519 | 0.00026302 | 0.00257981 | |
| From CR | | | | |
| | CR | GE | HU | PO |
| CR | | 1 | 1 | 1 |
| GE | 0 | | 0 | 0 |
| HU | 0 | 0 | | 0 |
| PO | 0 | 0 | 0 | |
| From GE | | | | |
| | CR | GE | HU | PO |
| CR | | 0 | 0 | 0 |
| GE | 1 | | 1 | 1 |
| HU | 0 | 0 | | 0 |
| PO | 0 | 0 | 0 | |
| From HU | | | | |
| | CR | GE | HU | PO |
| CR | | 0 | 0 | 0 |
| GE | 0 | | 0 | 0 |
| HU | 1 | 1 | | 1 |
| PO | 0 | 0 | 0 | |
| From PO | | | | |
| | CR | GE | HU | PO |
| CR | | 0 | 0 | 0 |
| GE | 0 | | 0 | 0 |
| HU | 0 | 0 | | 0 |
| PO | 1 | 1 | 1 | |
| To CR | | | | |
| | CR | GE | HU | PO |
| CR | | 0 | 0 | 0 |
| GE | 1 | | 0 | 0 |
| HU | 1 | 0 | | 0 |

| | | | | |
|---|---|---|---|---|
| PO | 1 | 0 | 0 | |
| To GE | | | | |
| | CR | GE | HU | PO |
| CR | | 1 | 0 | 0 |
| GE | 0 | | 0 | 0 |
| HU | 0 | 1 | | 0 |
| PO | 0 | 1 | 0 | |
| To HU | | | | |
| | CR | GE | HU | PO |
| CR | | 0 | 1 | 0 |
| GE | 0 | | 1 | 0 |
| HU | 0 | 0 | | 0 |
| PO | 0 | 0 | 1 | |
| To PO | | | | |
| | CR | GE | HU | PO |
| CR | | 0 | 0 | 1 |
| GE | 0 | | 0 | 1 |
| HU | 0 | 0 | | 1 |
| PO | 0 | 0 | 0 | |

Table S6. Information related to the genetic dataset used in this study

| Deme | Number of sequences | Date of the first sampled sequence | Date of the last sampled sequence |
|---|---|---|---|
| Poultry farms in Czech Republic | 36 | 2 January 2017 | 28 February 2017 |
| Poultry farms in Germany | 79 | 9 November 2016 | 9 May 2017 |
| Poultry farms in Hungary | 30 | 1 November 2016 | 20 April 2017 |
| Poultry farms in Poland | 45 | 1 December 2016 | 8 March 2017 |
| Wild birds in the four countries | 130 | 19 October 2016 | 11 March 2017 |

**Figures**

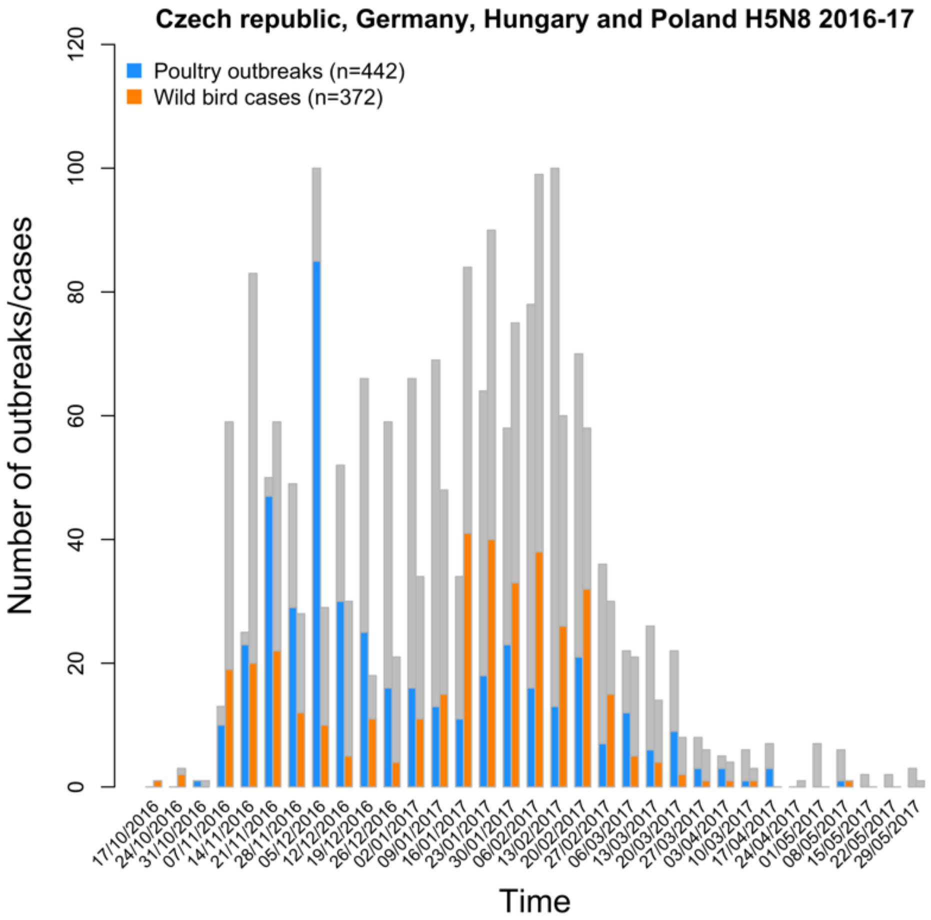


Figure S1. Temporal distribution of the number of poultry farm outbreaks and wild bird cases in Czech Republic, Germany, Hungary and Poland in comparison to the total number of outbreaks/cases reported in Europe (source: empres-i, 2017). According to this figure, the epidemic in the four countries can be described by four phases (two phases during which the number of outbreaks and cases was increasing: from October to November 2016 and January 2017, two phases during which the number of outbreaks and cases was decreasing: in December 2016 and from February to May 2017)

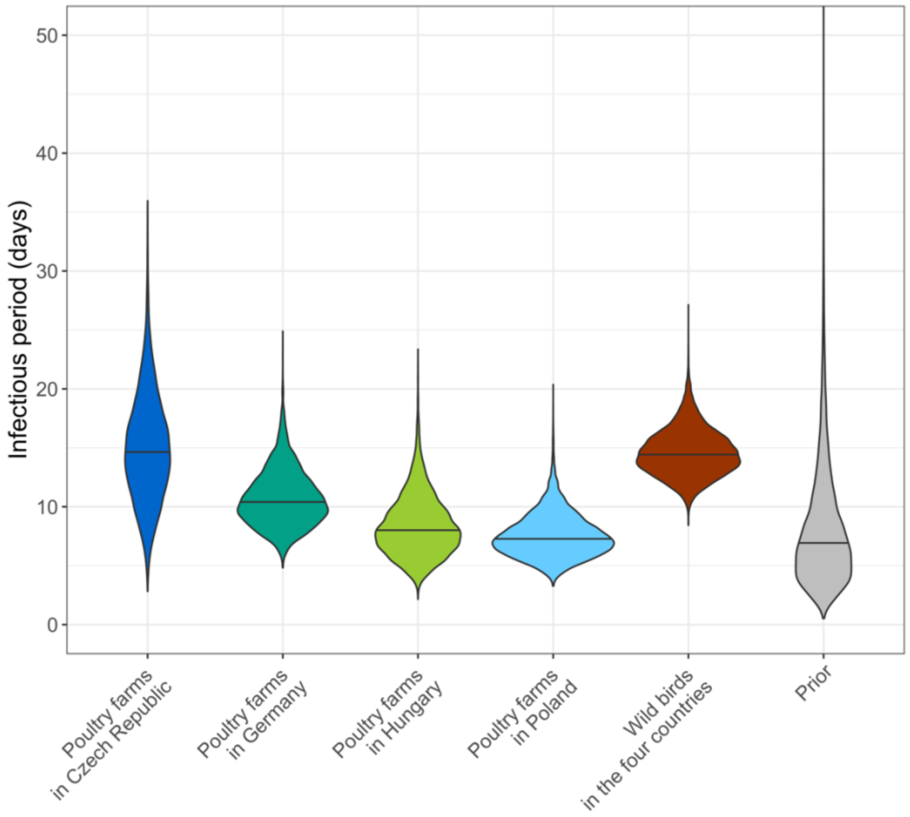


Figure S2. Posterior distributions for the infectious period (days) per deme. Solid horizontal lines represent median values.

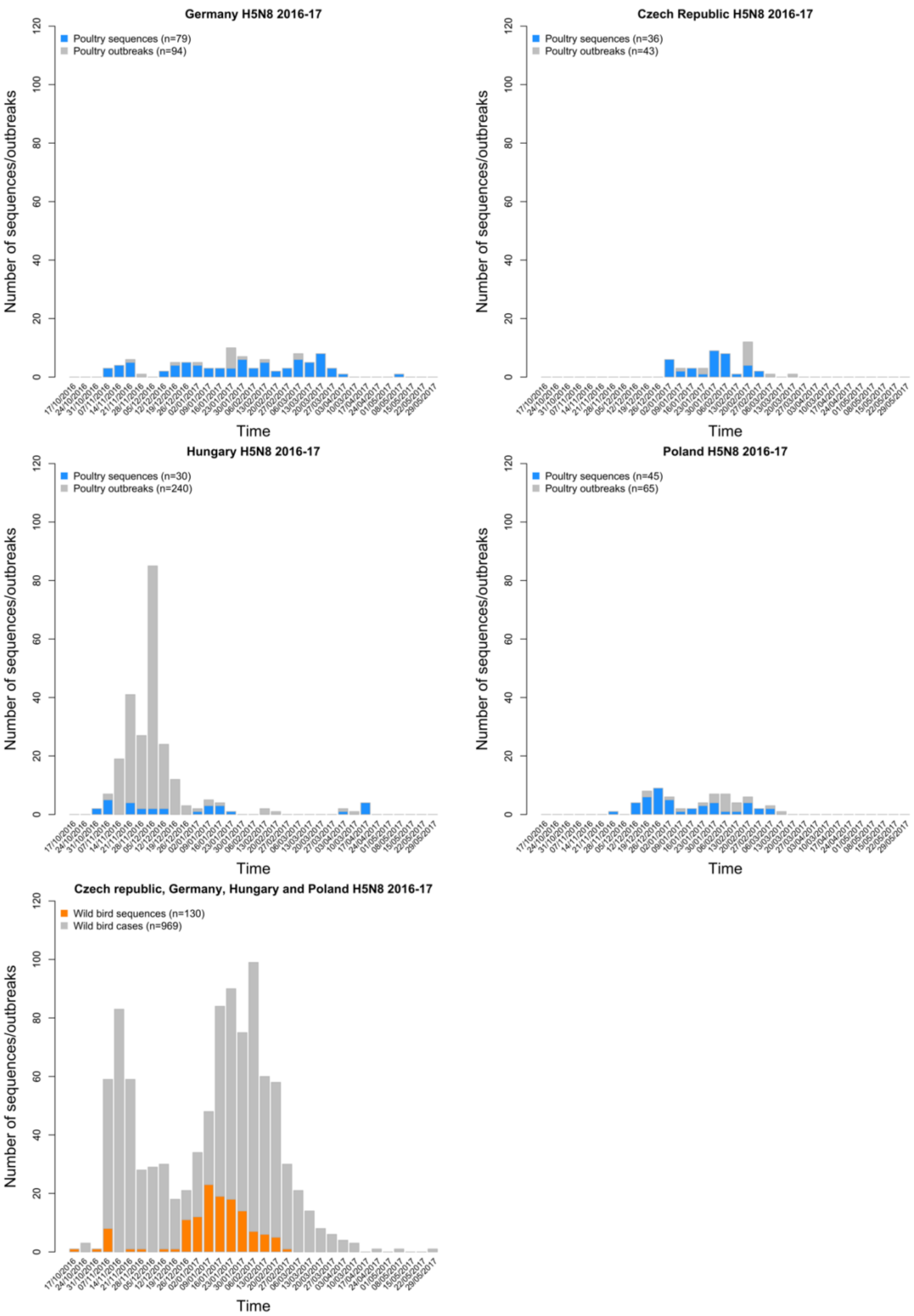


Figure S3. Temporal distribution of the number of H5N8 genetic sequences and outbreaks/cases per deme

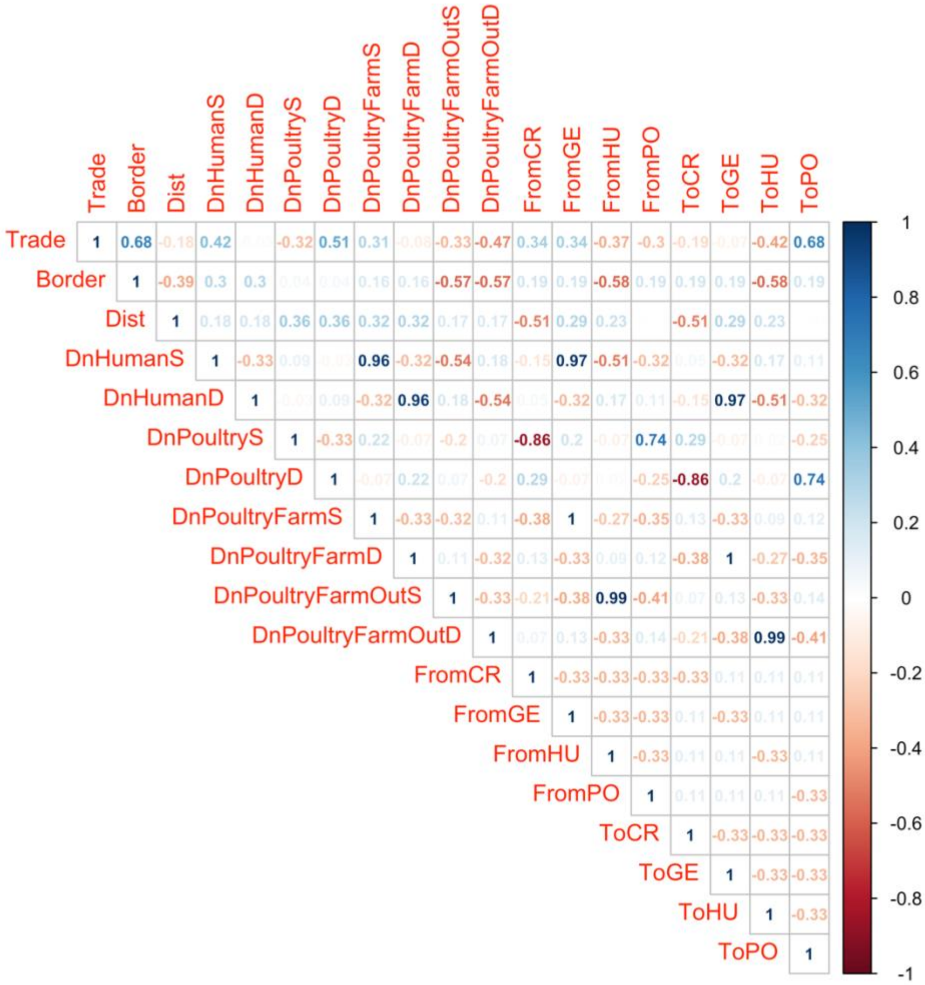


Figure S4. Correlation matrix of predictors. Upper half of the matrix shows the Pearson correlation coefficient. To avoid collinearity among predictors, predictors were removed when the Pearson correlation exceeded > 0.7. There were eight predictors included in the model: the live poultry trade, the poultry density in the source and destination deme, the poultry farm density in the source and destination deme, the farm outbreak density in the source and destination deme and the distance between countries' centroids

**References**


1. P. Alarcon, *et al.*, Comparison of 2016–17 and previous epizootics of highly pathogenic avian influenza H5 Guangdong lineage in Europe. *Emerg. Infect. Dis.* **24**, 2270 (2018).

2. A. Fusaro, *et al.*, Genetic Diversity of Highly Pathogenic Avian Influenza A(H5N8/H5N5) Viruses in Italy, 2016–17. *Emerg. Infect. Dis.* **23**, 1543–1547 (2017).

3. N. Beerens, *et al.*, Multiple Reassorted Viruses as Cause of Highly Pathogenic Avian Influenza A(H5N8) Virus Epidemic, the Netherlands, 2016. *Emerg. Infect. Dis.* **23**, 1974–1981 (2017).

4. D. A. Grear, J. S. Hall, R. J. Dusek, H. S. Ip, Inferring epidemiologic dynamics from viral evolution: 2014-2015 Eurasian/North American highly pathogenic avian influenza viruses exceed transmission threshold, R0 = 1, in wild birds and poultry in North America. *Evol. Appl.* **11**, 547–557 (2018).

5. A. Andronico, *et al.*, Highly pathogenic avian influenza H5N8 in south-west France 2016–2017: A modeling study of control strategies. *Epidemics* **28**, 100340 (2019).

6. I. Iglesias, *et al.*, Reproductive ratio for the local spread of highly pathogenic avian influenza in wild bird populations of Europe, 2005–2008. *Epidemiol. Infect.* **139**, 99–104 (2011).

7. K. Willgert, *et al.*, Transmission of highly pathogenic avian influenza in the nomadic free-grazing duck production system in Viet Nam. *Sci. Rep.* **10**, 1–11 (2020).

8. C. Leyson, *et al.*, Pathogenicity and genomic changes of a 2016 European H5N8 highly pathogenic avian influenza virus (clade 2.3. 4.4) in experimentally infected mallards and chickens. *Virology* **537**, 172–185 (2019).

9. S. Napp, N. Majó, R. Sánchez-Gónzalez, J. Vergara-Alert, Emergence and spread of highly pathogenic avian influenza A(H5N8) in Europe in 2016-2017. *Transbound. Emerg. Dis.* **65**, 1217–1226 (2018).

10. E. Świętoń, K. Śmietanka, Phylogenetic and molecular analysis of highly pathogenic avian influenza H5N8 and H5N5 viruses detected in Poland in 2016–2017. *Transbound. Emerg. Dis.* **65**, 1664–1670 (2018).

11. EFSA, *et al.*, Avian influenza overview October 2016–August 2017. *EFSA J.* **15**, e05018 (2017).

12. P. Lemey, *et al.*, Unifying viral genetics and human transportation data to predict the global transmission dynamics of human influenza H3N2. *PloS Pathog* **10**, e1003932 (2014).
